# Cryo-electron tomography of the herpesvirus procapsid reveals interactions of the portal with the scaffold and a shift on maturation

**DOI:** 10.1101/2020.09.30.320457

**Authors:** Michael H. C. Buch, William W. Newcomb, Dennis C. Winkler, Alasdair C. Steven, J. Bernard Heymann

## Abstract

Herpes simplex virus type 1 (HSV-1) requires seven proteins to package its genome through a vertex in its capsid, one of which is the portal protein, pUL6. The portal protein is also thought to facilitate assembly of the procapsid. While the portal has been visualized in mature capsids, we aimed to elucidate its role in the assembly and maturation of procapsids using cryo-electron tomography. We identified the portal vertex in individual procapsids, calculated a subtomogram average, and compared that with the portal vertex in empty mature capsids (A-capsids). The resulting maps show the portal on the interior surface with its narrower end facing outwards, while maintaining close contact with the capsid shell. In the procapsid, the portal is embedded in the underlying scaffold, suggesting that assembly involves a portal – scaffold complex. During maturation, the capsid shell angularizes with a corresponding outward movement of the vertices. We found that in A-capsids, the portal translocates further than the adjacent capsomers and strengthens its contacts with the capsid shell. Our methodology also allowed us to determine the number of portal vertices in each capsid, with most having one per capsid, but some none or two, and rarely three. The predominance of a single portal per capsid supports facilitation of the assembly of the procapsid.

**Importance:** Herpes simplex virus type 1 (HSV-1) infects a majority of humans, causing mostly mild disease but in some cases progressing towards life-threatening encephalitis. Understanding the life cycle of the virus is important to devise countermeasures. Production of the virion starts with the assembly of an icosahedral procapsid including DNA packaging proteins at a vertex, one of which is the dodecameric portal protein. The procapsid then undergoes maturation and DNA packaging through the portal, driven by a terminase complex. We used cryo-electron tomography to visualize the portal in procapsids and compare them to mature empty capsids. We found the portal located inside one vertex interacting with the scaffold protein in the procapsid. On maturation, the scaffold dissociates, the capsid angularizes, and the portal moves outward, associating closely with the capsid shell. These transformations may provide a basis for the development of drugs to prevent HSV-1 infections.

## Introduction

Members of the *Herpesviridae* family cause persistent infections in humans, ranging from cold sores on the lips and genitals to shingles and tumors. While individual members of this family exhibit differences in pathogenicity and structural details, reflected in their classification into three subfamilies, their overall architecture is largely conserved. The large genome of HSV-1, the prototypical representative of *Herpesviridae*, is accommodated in a correspondingly massive and complex capsid. During their elaborate life cycle (1), nascent capsids assemble in the nucleus as spherical precursor particles called procapsids (2–4), which conform to T=16 quasi-icosahedral symmetry and contain a spherical internal scaffold. The scaffold shell consists of the viral proteins pUL26.5 and pUL26, of which the latter has an N-terminally module, the viral protease (4). Upon completion of the procapsid, the protease is activated and proceeds to cleave the scaffold, initiating maturation of the capsid. Maturation involves: (I) angularization of the capsid; (II) reorganization of the interactions that engage the triplex proteins as well as rearrangement of capsomers (1–3) from skew to more symmetric orientations; (III) conformational changes (2, 5–8) that establish interactions between capsomer subunits; and (IV) decoration of the mature capsid with additional proteins (9, 10). Maturation increases the capsid’s stability dramatically, countering the internal pressure exerted by the tightly packed genome (11). This process yields three capsid species, two of which remain in the nucleus. The A-capsid has a mature shell that does not contain any DNA and is essentially empty, while B-capsids retain a shrunken and somewhat disordered scaffold despite having matured into an angular shape. Only mature, DNA-filled capsids, called C-capsids, leave the nucleus and do so via the perinuclear space (reviewed in (12, 13)), to undergo the subsequent steps in the viral life cycle.

DNA packaging occurs at a capsid vertex and requires seven proteins, pUL6, pUL15, pUL17, pUL25, pUL28, pUL32 and pUL33 (14). pUL6 forms a circular dodecamer similar to the portals of bacteriophages (15) and may facilitate procapsid assembly (16). Heterotrimers of pUL17 (one copy) and pUL25 (two copies) are bound to the pentons and peripentonal triplexes (17–19). A terminase complex composed of pUL15, pUL28 and pUL33 provides the force for packaging and subsequent cleavage of the DNA (20–26). The final protein, pUL32, has not been found as a component of capsids (14). These proteins replace a penton presumably at one vertex of the capsid (27). While the structural disposition and transformations of some of these components have been elucidated, many details of assembly, maturation and packaging remain to be described.

Early attempts to locate the portal-bearing vertex in capsids employed cryo-electron tomography (cryoET) (28, 29). The tomograms were of low resolution (55-60 Å), complicating analysis. This led to placing the portal either at the level of capsomers (28) or inside the capsid shell (29). Subsequent analysis employing cryo-electron microscopy (cryoEM) and single particle analysis (SPA), indicated that the portal is located inside the capsid shell (30, 31). One of the difficulties in the SPA approach is that the overlap of one vertex with the large mass of the capsid (~100 fold ratio) makes it hard to distinguish a portal vertex from the penton vertices. Nevertheless, the extensive averaging of structural information means that correct assignments would dominate incorrect assignments distributed over the eleven other possibilities and reveal the location of the portal. The recent technological advances in cryoEM enabled the high resolution reconstruction of various herpesvirus capsids showing the portal on the inside of one vertex (18, 31–34).

One omission from the existing studies is the predisposition of the portal in the initial assembly product, the procapsid. It is also not clear that only one vertex bears a portal as assembly can occur without portal (2). In this study, we returned to cryoET to examine the portal vertices in procapsids and compared them to A-capsids as a control. In this effort, we developed the computational capability to find the portal vertices in a more reliable manner, allowing an estimate of the number of portal vertices in each capsid. From subtomogram averages of the portal vertices, we found abundant contacts between the portal and the scaffolding layer in the procapsid. We also found that the portal moves closer to the capsid shell on maturation.

## Results

### Production of procapsids

Procapsids were produced using the M100 mutant that lacks the viral protease VP24 required for rapid maturation (35). We followed a gentle purification protocol (5, 7) to minimize damage to the particles. For successful identification of the portal vertex in procapsids, several precautions were taken: since procapsids are fragile and malleable, we used an antibody binding to the major capsid protein (2) that stabilized the particles. We performed cryoET on the isolated procapsids, collecting and analyzing 80 tilt series where the particles were well separated (Fig. 1A).

**Fig. 1.**
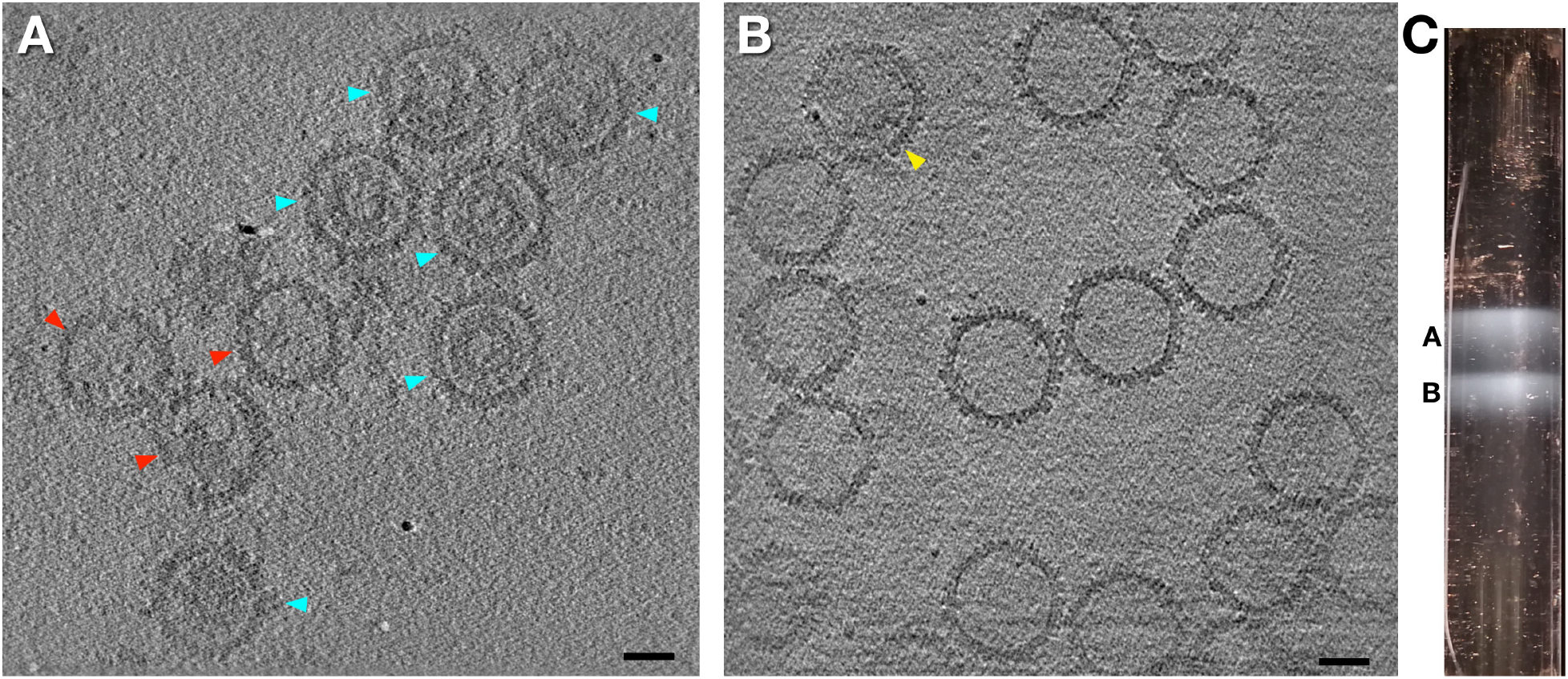
Isolated capsids. **A.** Central slice of a tomogram of procapsids. Some particles contained visibly degraded scaffold or showed imperfections (red arrowheads) and were excluded from further processing. We took care to include only particles that were separated, spherical, and had a visibly intact scaffold (turquoise arrowheads). **B.** Central slice of a tomogram of A-capsids. The individual particles are well defined and separated. Occasionally, a capsid with some internal content – probably some scaffold – were found in the field of view (yellow arrowhead). These particles were not included in further processing. **C.** Small-scale sucrose gradient of a capsid mix obtained during initial purification from a UL25-Null mutant, showing bands of A- and B-capsids. Scale bars, 500 Å.

### Production of capsid samples using a pUL25-deficient mutant

We decided to analyze mature A-capsids as a control to avoid interference from internal density features such as scaffolding (cf. B-capsids) or packaged DNA (cf. C-capsids). The established protocol for purifying HSV-1 nucleocapsids from a wild-type infection yields only a minor fraction of A-capsids. To increase their yield, we used a mutant lacking the vertex protein, pUL25, that is defective in DNA packaging (36). This virus replicates in a complementary African Green Monkey cell line transfected with the *UL25* gene, but in wild-type cells cannot produce progeny. In our protocol, centrifugation on small-scale sucrose gradients yielded bands for A-capsids and B-capsids but no band for the DNA-filled C-capsids (Fig. 1C). We isolated the UL25-Null A-capsid band (hereafter referred to as A-capsids) and collected tilt series for cryoET. Individual capsids in the tomograms were well spread out in the sample, and almost all capsids could be confirmed as A-capsids from their empty appearance (Fig. 1B).

### Identification of the portal vertex in cryo-electron tomograms

Our aim was to locate the portal using cryoET to take advantage of the straightforward identification of unique features in 3D maps. The approach used in an earlier tomographic protocol relied on simple template matching to find the portal vertex (28). Another protocol made use of the comparison of opposite vertices and assessing the density differences between vertices (30). In an effort to improve the detection of portal vertices, we developed a different algorithm aimed at detecting the vertex that is most different from the other vertices. We implemented this algorithm in a new program called bico added to the existing Bsoft package (37). An overview of our workflow is given in Fig. 2. From reconstructed tomograms, subtomogram volumes were extracted and aligned to an icosahedral reference, in our case a reconstruction of a HSV-1 C-capsid (accession number EMDB-7472 (38)).

**Fig. 2.**
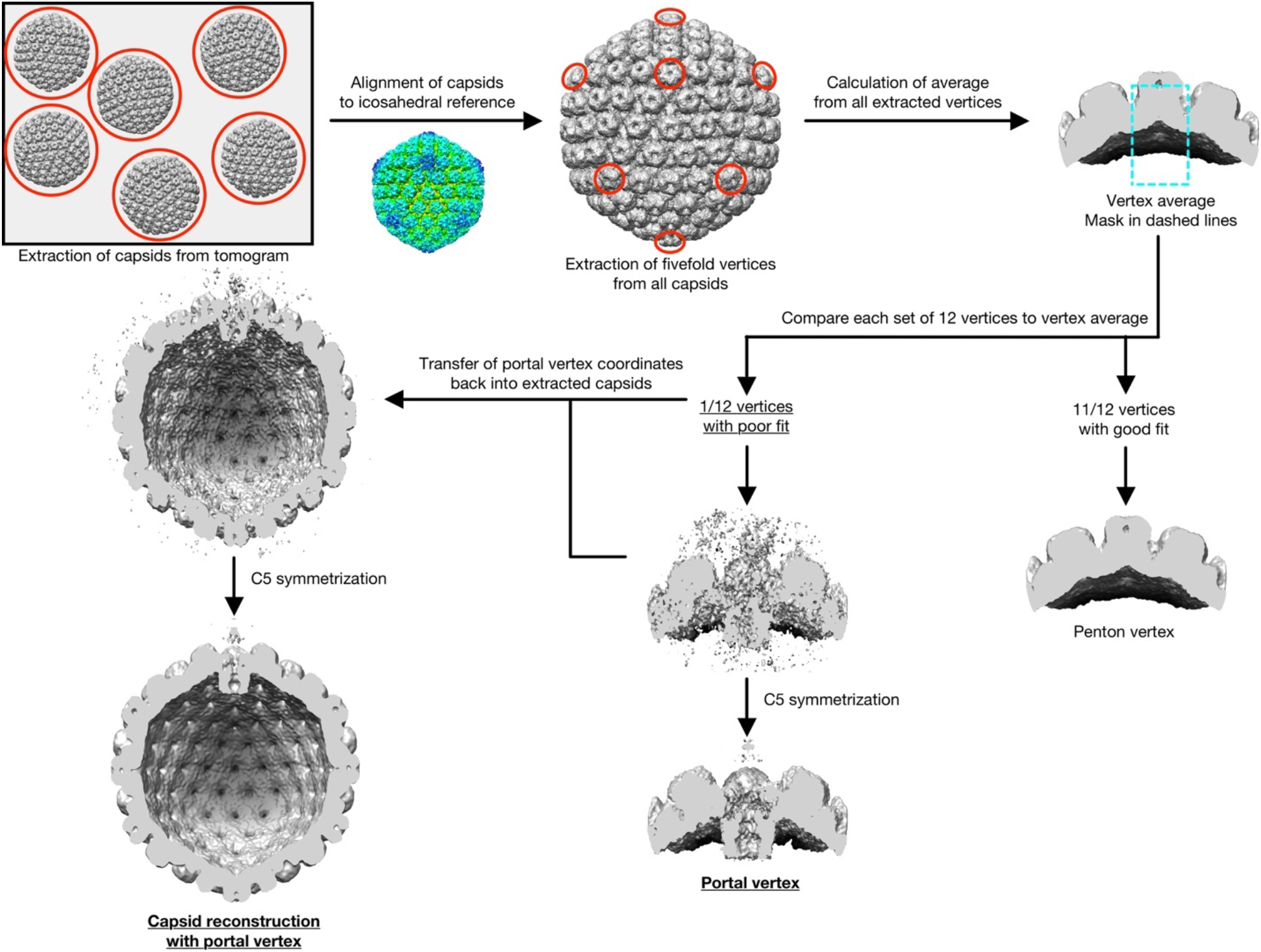
Workflow used to identify and average the portal vertices. Individual particles (volumes each containing a capsid) extracted from tomograms and aligned with an icosahedral reference using a frequency space mask to compensate for the missing wedge. This allows the location of the twelve 5-fold vertices of each capsid, which are then extracted and averaged. Each vertex is correlated to this average using a missing wedge mask in frequency space and a cylindrical real space soft mask to isolate the region in a vertex occupied by the penton or portal (cyan dashed line). The vertex with the lowest (worst) correlation is assumed to contain the portal. This is followed by a further refinement to find the best 5-fold orientation around the vertex axis. The coordinates of the portal vertices were then used to transform their corresponding capsids into orientations that placed the portal vertex on the z-axis at the top of the volume.

Next, the fivefold vertices of each capsid were extracted, and an average of all extracted and correctly oriented vertices was calculated. To focus only on the central part of a vertex where we expect the biggest difference between penton and portal, we used a soft cylindrical mask in the comparisons of each vertex with the average (see Materials and Methods for mask details). For each set of 12 vertices, bico then classified each vertex set into two subsets: the first subset contained the eleven best-fitting vertices. The second subset contained the vertex with the worst fit, taken to be the portal vertex. The coordinates of the identified portal vertices were then transferred back into the extracted and aligned capsids, and a reconstruction of the capsid featuring the portal vertex was calculated. To reduce noise, this reconstruction was symmetrized fivefold in the final step and filtered to 30 Å (just beyond the 34 Å resolution estimated by Fourier shell correlation – see Fig. S1).

### The portal-bearing vertex of UL25-Null A-capsids

We extracted a total of 164 subtomogram volumes of A-capsids from our tomograms. We identified the likely portal-bearing vertex in the map of each of the A-capsids and calculated an average (Fig. 3), obtaining a resolution of 34 Å (Fig. S1). In line with recent publications (18, 32, 33), we visualized the portal on the inside of the capsid, with its wider end facing towards the center of the particle (Fig. 3A). A superposed atomic model of the HSV-1 portal (PBD ID 6OD7 (32)) fits well into the portal density (Fig. 3B). The portal has a narrow end called the “clip domain” that engages the triplexes (green arrow in Fig. 3B) (for the assignment of this density as a triplex, see Huet et al. (31)). The widest part of the portal, the “wing domain” extends towards the floors of the neighboring capsomers (blue arrow in Fig. 3B). The overall appearance and placement of this connection is consistent with a set of β-barrels and spine helices originating from the pUL19 hexons that were identified earlier in a high-resolution reconstruction (32). Knowing what it looks like, we could identify the portal vertex in individual particles in our tomograms (white arrows in Fig. 3C).

**Fig. 3.**
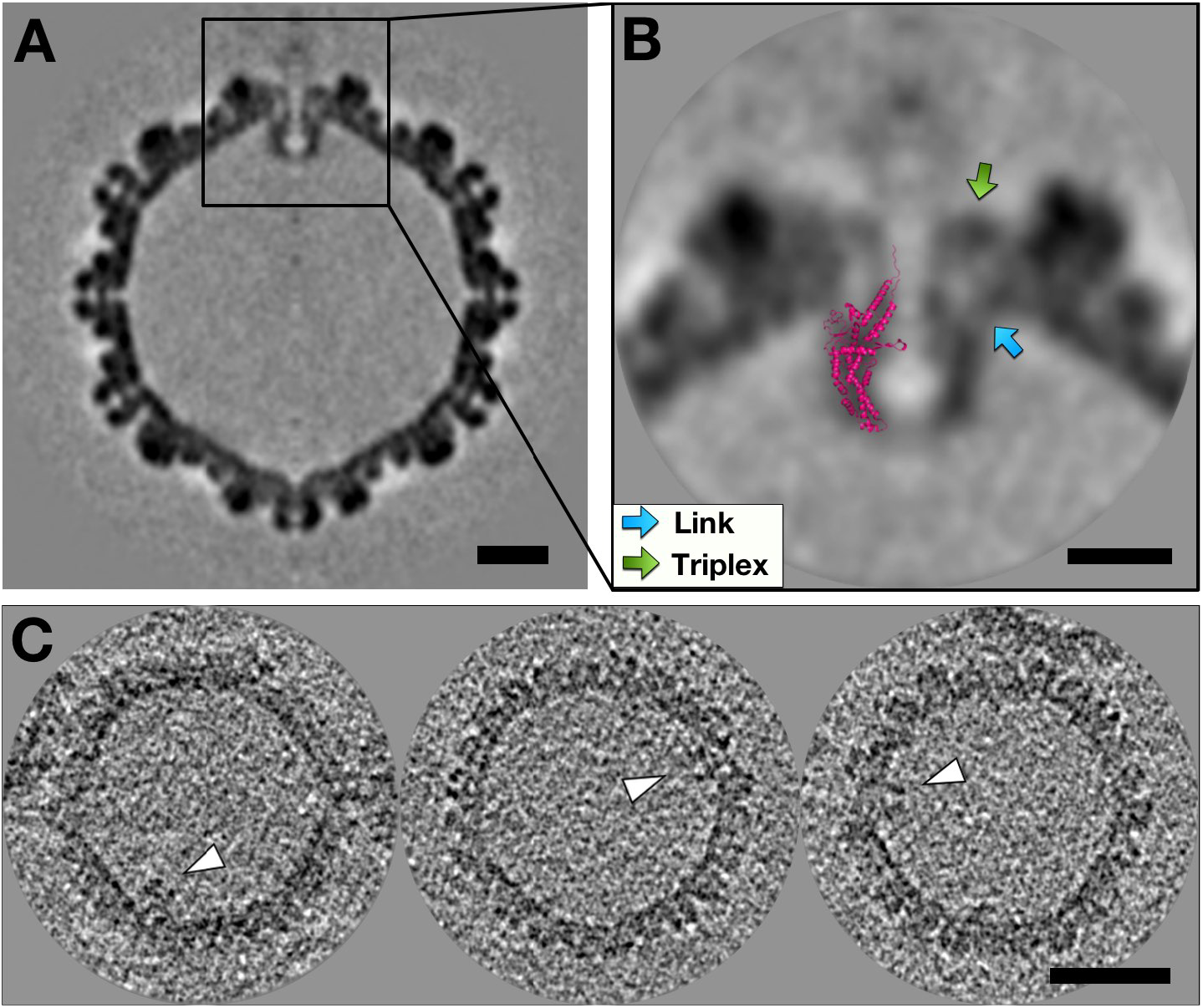
The portal vertex in UL25-Null A-capsids. **A.** Central slice of the C5-symmetrized capsid with the portal vertex located at the top. Scale bar, 200 Å. **B.** Magnified view of the vertex. The portal is located beneath the capsid floor with its wider end facing the inside of the particle. A high-resolution model (pink, PDB ID 6OD7 (32)) is superimposed on the left half of the portal density. The portal forms connections to the triplex (green arrow) and the capsid floor (blue arrow indicates a link). Note that there is a diffuse density above the portal vertex that could be residual terminase. Scale bar, 100 Å. **C.** Slices from tomograms of A-capsids that traverse the portal in each (white arrowheads). Scale bar, 500 Å.

### The portal in procapsids interacts with the scaffold

We selected tomograms with well-separated spherical particles (Fig. 1A) and extracted 368 subvolumes. As with A-capsids, we identified the portal-bearing vertex, taking care to adjust the mask so that it excluded the scaffold, which might otherwise outweigh the portal density during vertex sorting. After identifying the portal vertex in each procapsid, we calculated an average (Fig. 4) with a resolution of 34 Å (Fig. S1). Consistent with the A-capsids, the portal is located inside the capsid shell (Fig. 4A), and again agrees with an atomic portal model (Fig. 4B). The portal penetrates deeply into the scaffold layer (yellow arrow in Fig. 4B), almost to the widest part of its wing, a position that allows the formation of extensive interactions. Interestingly, the nominally spherical scaffold shows a degree of perturbation caused by the portal, giving the scaffold an overall teardrop shape (Fig. 4A). We also note that – apart from in the portal region - the scaffold shell shows up in slices as three concentric shells (arrows in Fig. 4A) consistent with previous studies (5, 7). Only tenuous contacts (blue arrow in Fig. 4B) are formed between the portal wing domain and the surrounding hexons, but the clip domain appears to interact with the triplexes (green arrow in Fig. 4B). The vertex overall has a funnel-like shape that likely accommodates DNA insertion by the terminase complex. Finally, the procapsid hexons also display faint residual density on their exterior parts that we attribute to the 6F10 antibody used to stabilize the procapsids during isolation (Fig. 4A, 5C).

**Fig. 4.**
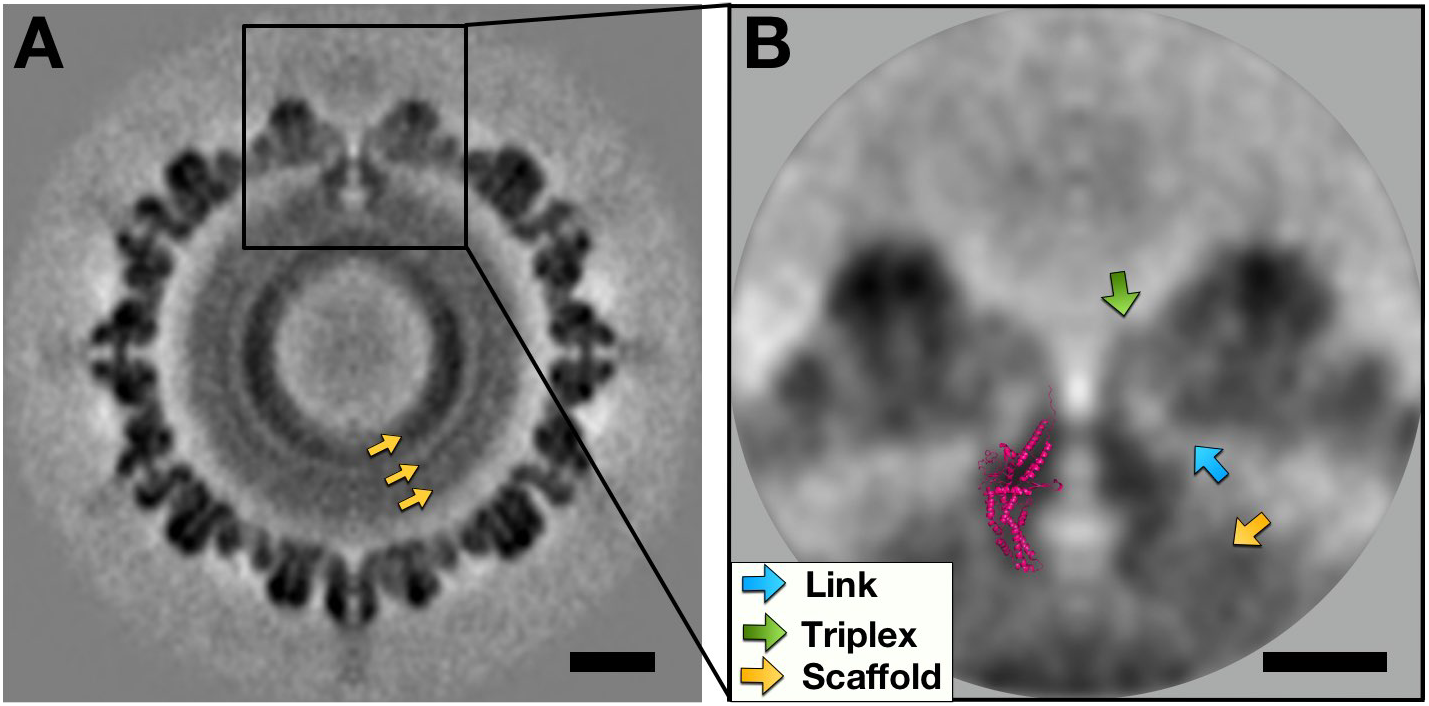
The portal vertex in procapsids. **A.** Central slice of the C5-symmetrized procapsid with the portal vertex at the top, showing distinct rings of scaffold density (yellow arrows). Scale bar, 200 Å. **B.** Magnified view of the portal vertex with a superimposed atomic model (pink, PDB ID 6OD7 (32)). The portal is mainly connected to the capsid through the triplex (green arrow) with a thin connection to the capsid floor (blue arrow). The bottom of the portal is embedded in the scaffold (yellow arrow). Scale bar, 100 Å.

### Conformational changes and movement of the portal during maturation

While the portal orientation is similar in procapsids and A-capsids, contrasts in vertex configuration come to light in a side-by-side comparison (Fig. 5). Upon maturation, the capsid becomes angular, causing all the vertices (both penton and portal) to translocate outwards. We assessed the movement of the pentons by shifting the penton vertex of the procapsid outwards until it overlays the A capsid penton, yielding a translation of 29 Å. Similarly, we measured the translation of the portal to be 54 Å, thus moving closer to the capsid shell. The portal in Fig. 5A appears to be somewhat separated from the capsid shell but is much closer in the mature capsid (Fig. 5B). During maturation, the capsid shell stabilizes by closure of the floor and burying interfaces between the capsomers (8). Our results indicate that the portal also adopts a more stable interaction with the mature capsid shell.

**Fig. 5.**
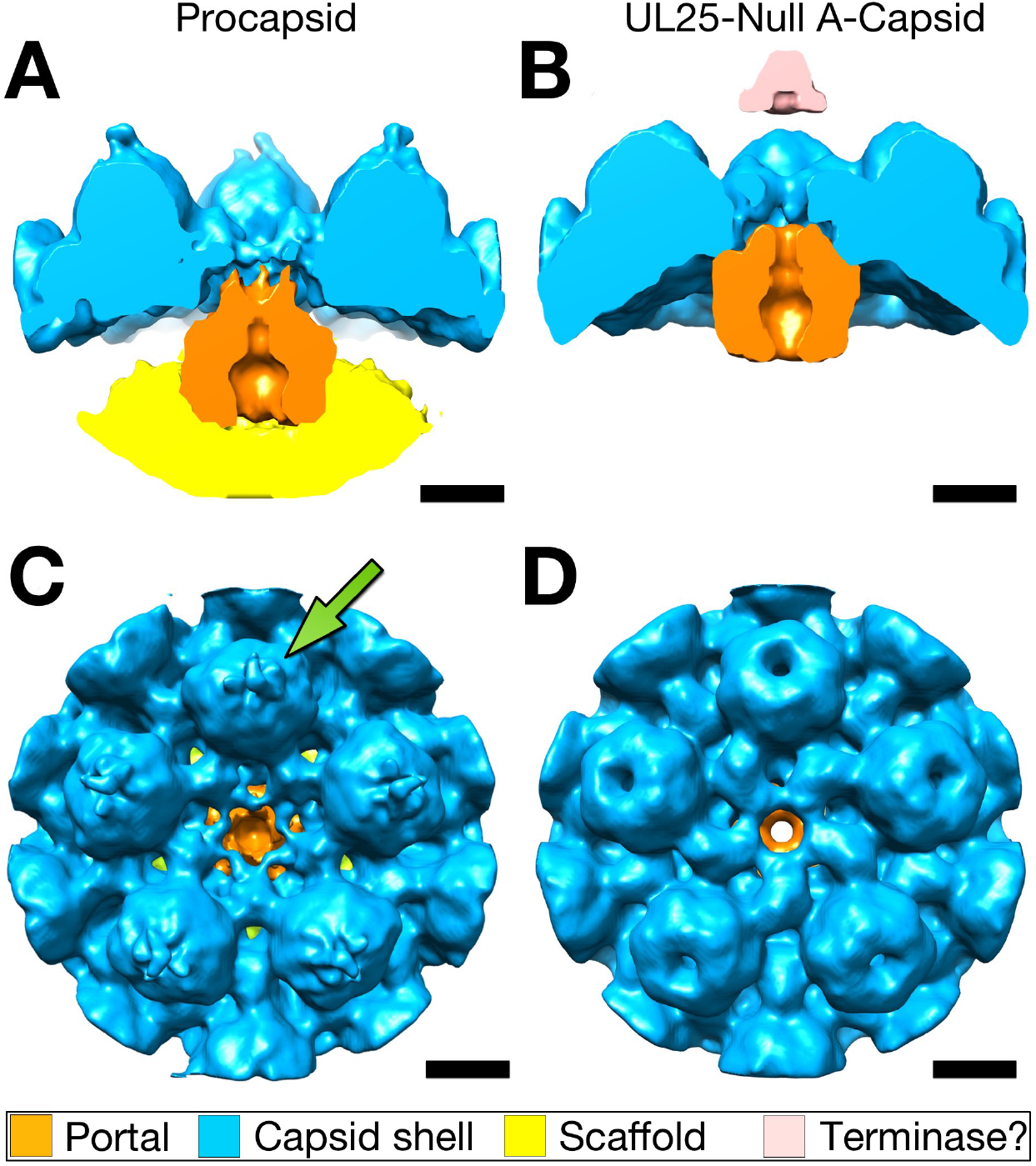
Comparison of portal vertices in procapsid and UL25-Null A-capsid. **A.** Side view cut through the procapsid portal vertex, showing the portal (orange) connecting to the capsid shell (blue) and embedded in the scaffold (yellow). **B.** Side view of the mature A-capsid portal vertex, with the portal (orange) very closely associated with the capsid shell (blue). The diffuse density above the vertex (pink) may be residual terminase. **C.** Top view of the procapsid portal vertex with faint density associated with the tops of the hexons attributed to the 6F10 antibody used in the preparation of the particles (green arrow). **D.** Top view of the A-capsid portal vertex. Upon maturation, the triplexes surrounding the vertex shift and constrict the vertex channel, and the hexons transform from skewed ovals towards hexagonal symmetry. Scale bars, 100 Å.

In addition, we observe a shift in the triplex network surrounding the portal vertex, which narrows the vertex channel. These changes become even more apparent when the vertices are viewed from the top (Fig. 5C,D). While triplexes are somewhat weakly defined in the procapsid portal vertex, they become much more visible in the A-capsid and even extend towards the fivefold symmetry axis. It is possible that some of this extending density could be ascribed to components of the tegument, particularly pUL17. We recall that, upon maturation, the hexons rearrange from a skewed orientation to true hexameric symmetry (8).

We also compared the triplexes of the portal vertex with those of the opposing penton vertex in both maps. The triplexes in the procapsid portal vertex are poorly defined (Fig. 6A), making it hard to ascertain whether they are in the same conformation as those at the opposing penton vertex (Fig. 6B). In contrast, the orientations of the triplexes in the A-capsid are clearly the same for both vertices (Fig. 6C,D).

**Fig. 6.**
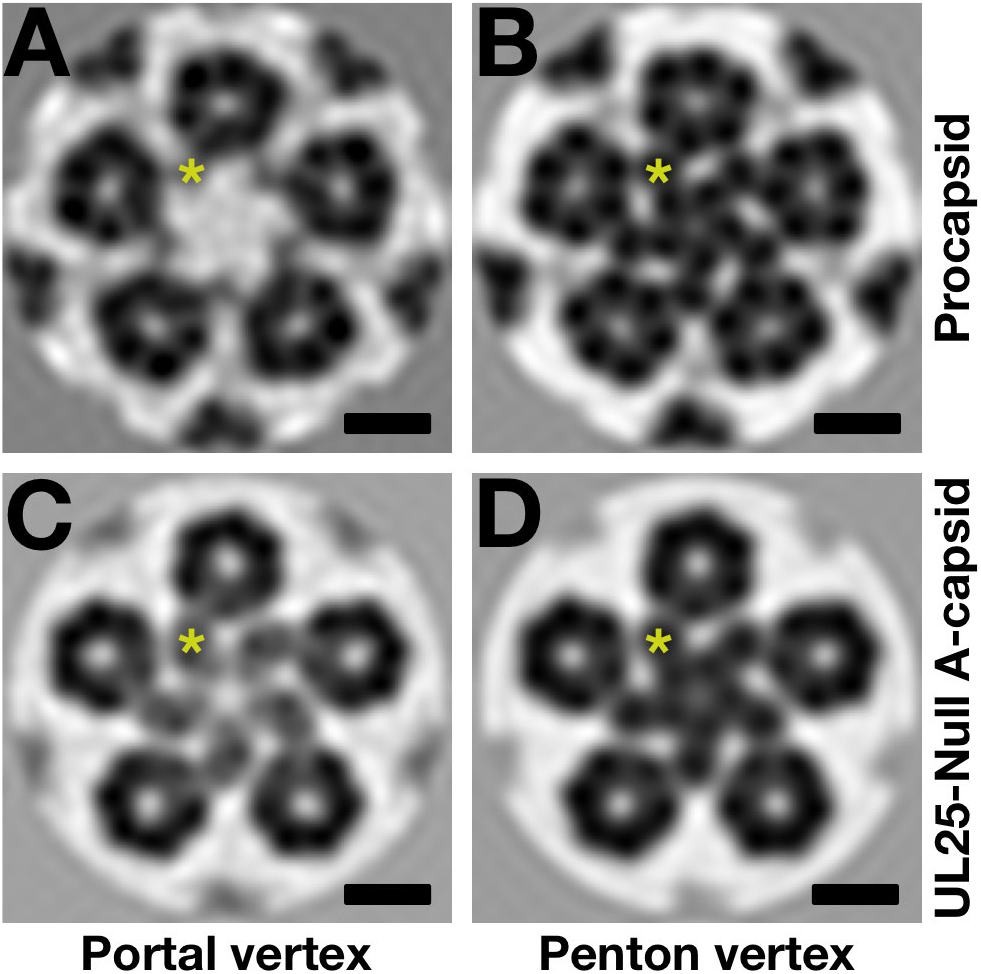
The arrangement of triplexes at the vertices. The triplexes (yellow stars) at the procapsid portal vertex (**A**) are recessed relative to those at the opposite penton vertex (**B**) (slices at a radius of 307 Å). In contrast, the triplexes at the A-capsid portal vertex (**C**) are closer in conformation to those at the opposite penton vertex (**D**) (slices at a radius of 325 Å). Note that the vertices in the procapsid move outwards by ~29 Å on maturation, accounting for the differences in slices shown here. Scale bars, 100 Å.

### The number of portal vertices per capsid

While we implicitly assumed there is exactly one portal per capsid, that is not necessarily true. To determine the number of portal vertices in each capsid, we examined the distribution of correlation coefficients for the 12 vertices. Our aim was to generate two clusters: the few with the worst correlation coefficients indicating portal vertices, and the rest as penton vertices. To separate the clusters, we took the biggest difference between successive correlation coefficients in a sorted list as the distinction between penton and portal vertices. Because we expected only a few portal vertices, only the six with the lowest coefficients were assessed. However, this strategy assumes there is at least one portal vertex. We compared the difference between the two worst vertices with the next three differences and assumed that when it is the lowest, it indicated a capsid without a portal vertex. The resultant distributions are shown in Fig. 7, giving average portal vertices per capsid of 1.07 for procapsids and 1.25 for A-capsids, consistent with previous estimations (16, 27). In our subtomogram averages, we included all of the capsids, but we do not consider the minor contribution of the capsids without portals to significantly change our conclusions. About 65 % of the capsids have a single portal vertex, in agreement with the 73 % reported for A-capsids (28). A still significant number (~20 %) have two portal vertices, while those with three are rare. If the incorporation of portal vertices follows a random pattern, the expected distribution would be binomial in nature (blue lines in Fig. 7). The predominance of a singly incorporated portal indicates that the assembly is more specific than a random process.

**Fig. 7.**
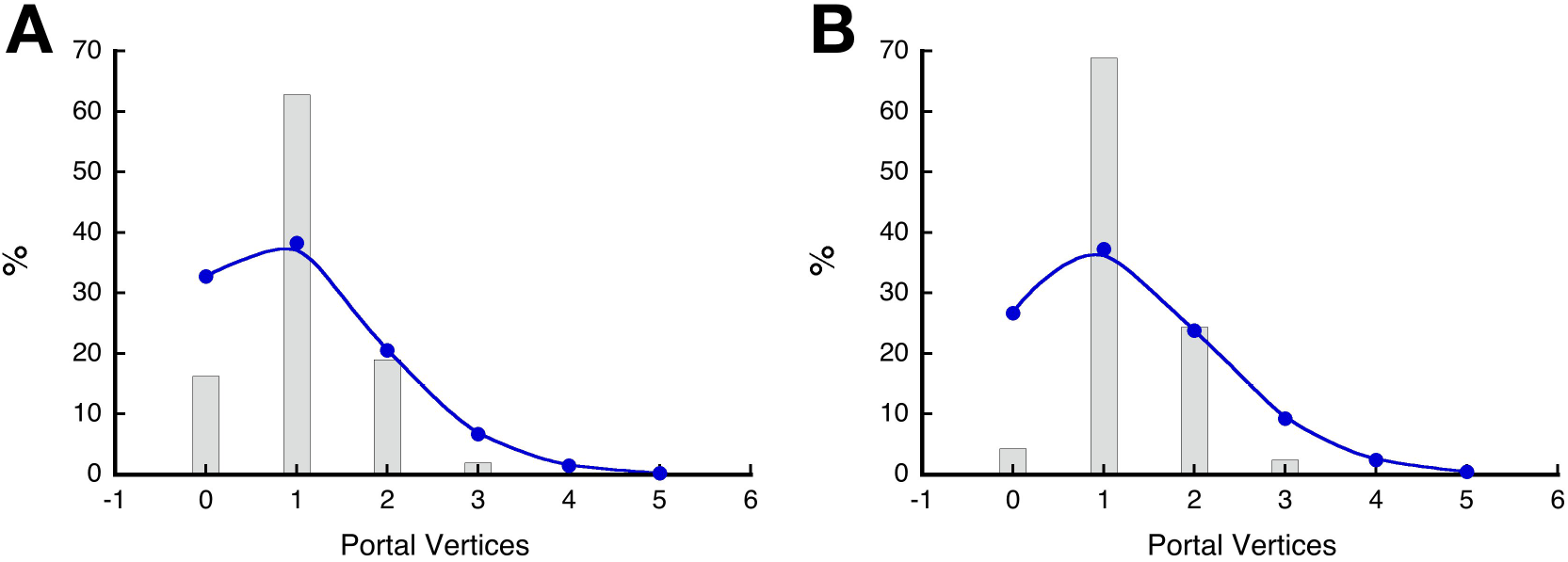
Estimating the number of portal vertices (bars) per procapsid (A) an A-capsid (B). The blue line in each plot is a binomial distribution calculated from the average number of portal vertices per capsid as an indication of expected random incorporation.

## Discussion

### Localization and interactions of the portal in procapsids compared to A-capsids

In a critical step of the HSV-1 life cycle, newly synthesized DNA is threaded into the capsid which undergoes extensive conformational changes as it matures (5, 8) although the latter process can occur without DNA packaging (8) and may precede DNA packaging *in vivo* (39). In both herpesviruses and tailed bacteriophages, DNA is inserted through a fivefold vertex via the dodecameric portal protein and driven by terminase complex. While the overall structure of the HSV-1 portal, pUL6, has been known for some time (15), its orientation and placement in the context of the capsid shell have been subject to debate. The portal’s overall shape makes it difficult to distinguish visually from capsomer pentons in micrographs, making consistent alignment of 2D projections a challenge. In an earlier study, we used cryoET, aiming to characterize the portal vertex of HSV-1 (28). Here, we have refined this approach by developing a new program designed to identify the portal vertex on individual capsids, thereby allowing coherent averaging. Applying it to HSV-1 procapsids and A-capsids, we located the portal on the inside of the particle, consistent with several recent SPA-based studies (18, 30, 32–34). In confirmation, an atomic model of pUL6 (32) fits well into our portal density (Fig. 8). This density extends beyond the clip domain of the model, presumably representing some of residues 308-515, which were not modeled (corresponding residues are also unmodeled in a reconstruction of the Epstein-Barr-virus portal (40)). Interestingly, a corresponding turret-shaped density was discovered in the portal vertex of Kaposi’s Sarcoma Herpesvirus (KSHV) (34); it was theorized that this helical turret is an extension of the portal’s clip domain and could be a docking site for the terminase complex. If such a component is indeed an additional protein, it may only be added to mature particles upon the successful retention of DNA.

**Fig 8.**
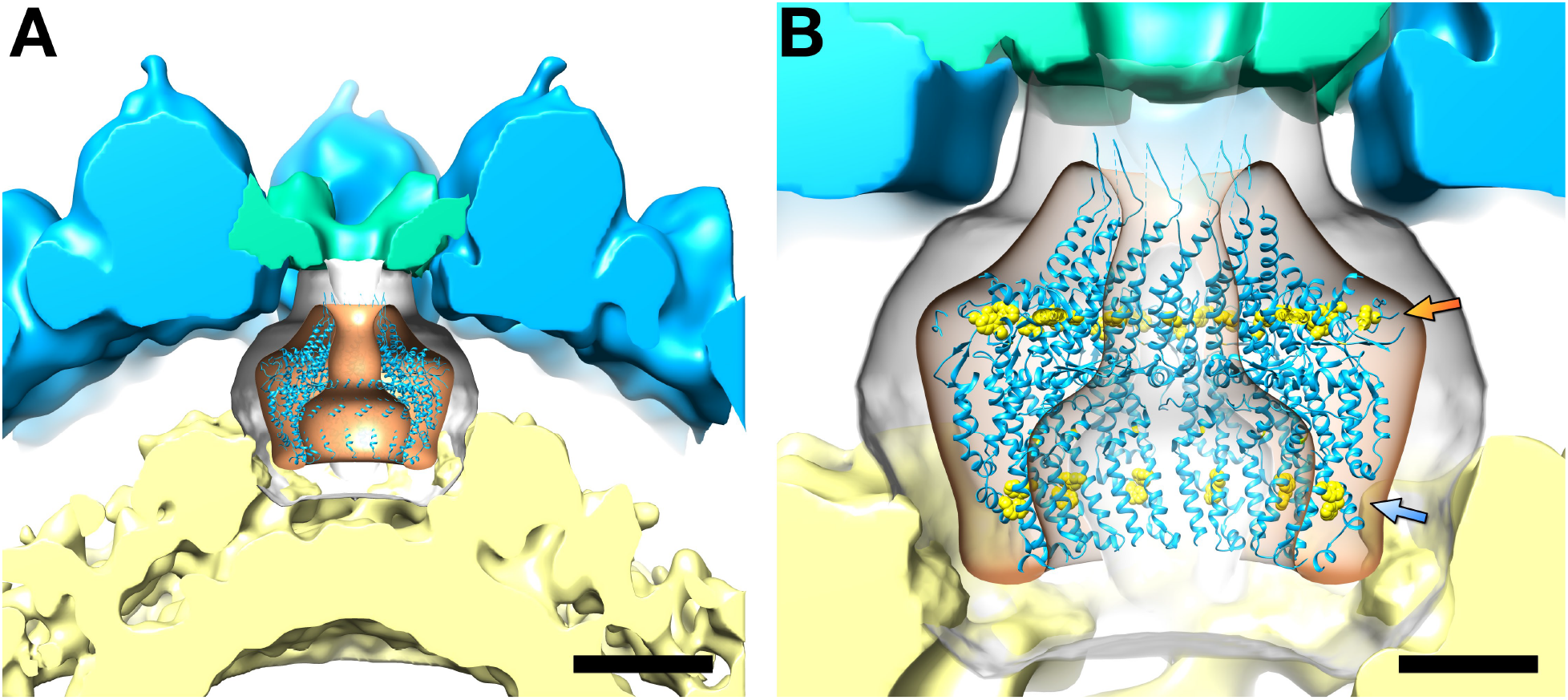
The model of the portal within the context of the procapsid portal vertex. **A.** The portal model (PDB ID 6OD7 (32)) fitted into the portal density (orange) within a lower threshold envelope (grey) to show the connections to the triplexes around the portal (green) and the scaffold (yellow). The connections to the capsid shell (composed of the major capsid protein and other triplexes shown in blue) are not resolved in this map. Scale bar, 100 Å. **B.** Close-up view of the portal with key tryptophan residues (yellow balls) at the wing (orange arrow: W163, W241, W262, W532) and at the bottom (blue arrow: W90, W127) that were shown to interact with the scaffold (41). Scale bar, 40 Å.

Applying our workflow to procapsids led to several novel observations. The procapsid portal is located inside the vertex forming contacts with the scaffolding shell and deforming the scaffold proteins’ spherical arrangement at the interaction site. Conserved tryptophan residues have been identified in HSV-1 pUL6 ((41), see also Fig. 2 in (42)) that play crucial roles in interaction between portal and, most likely, conserved scaffold residues 143-151 of pUL26, particularly YYPGE (43, 44). Figure 8B shows these critical tryptophan residues arrange in a ring in the wing domain (orange arrow in Fig. 8B) as well as a ring in the base of the portal (blue arrow in Fig. 8B). We know the scaffold C-terminus is bound to the capsid (45) which puts it in the proximity of the portal wing domain. Conserved tryptophan residues are a feature not only of HSV-1 but also of some bacteriophages ((46), reviewed in (42)). The scaffold probably shares the symmetry of the surface shell with which it co-assembles but is not highly ordered and its apparent sphericity is enhanced by averaging. From previous studies we know that the protease at the N-terminal part of about 10 % of the scaffold proteins is located towards the center of the procapsid (4) and the C-terminus interacts with the capsid shell (47) and portal (43). This means the protein is in an extended radial conformation consistent with a side-by-side packing of coiled coil dimers as inferred by Pelletier et al. (48). In the procapsid we see three peaks in the concentric density shells (Fig. 4) in agreement with the three domains proposed by Trus et al. (7).

### Incorporation of the portal during procapsid assembly

The stage at which the portal is incorporated into the procapsid is still unknown, with possibilities that it could initiate assembly, inserted at some intermediate point, or at the end (16). Procapsids can assemble *in vitro* without portal present (2, 3) and some of the capsids we analyzed appear to lack a portal vertex (Figure 7). We also found an average copy number slightly larger than one and consistent with previous studies (16, 27). If one assumes a random model of incorporation, it is expected to follow a binomial distribution. However, Figure 7 shows it clearly deviates from a binomial pattern, suggesting a predisposition towards a single portal vertex per capsid. This suggests that the portal may facilitate procapsid assembly and thus ensure that a majority has at least one portal vertex. However, if it initiates assembly, it does not preclude incorporation of one or two additional portals.

### Morphological changes upon capsid maturation

Cleavage of the C-terminal 25 residues of the scaffolding protein by the protease (47) triggers maturation of the capsid shell. Our preparation of procapsids is defective in the protease to allow us to purify it. Nevertheless, the capsids are still able to mature slowly (8). On cleavage, the scaffold changes from extended in the procapsid (i.e., large core particles) to somewhat contracted in the B-capsid (i.e., small core particles). At the same time, the vertices of the capsid shell shift outwards during angularization by ~29 Å, with the portal itself moving even further by ~54 Å (note that the capsid shells in A- and B-capsids are very similar (31)). The portal therefore disengages from the scaffold to some extent, while strengthening interactions with the capsid shell. The portal movement is accompanied by the establishment of a support network with the newly established capsid floor (termed suspension network in another study (32)).

The overall shape of the portal vertex differs significantly between procapsids and mature capsids (Fig. 6). In procapsids, we note that the center of the vertex is essentially open, which would, in a biological context, be advantageous for the insertion of DNA. Triplex density is present around the portal vertex, but it is somewhat fainter than around a penton vertex, which could result from a lack of interaction between the portal and the procapsid itself. In the mature capsid, the vertex is narrower, and it appears that density originating at the triplexes extends towards the vertex axis. This extended density may indeed be contributed by components of the tegument, particularly pUL17. While our A-capsids lack pUL25, it has been shown that procapsids contain practically no pUL25 (25), and given that the procapsid undergoes substantial conformational changes during maturation, it is likely that no tegument component protein is present on these immature particles. The extended, star-shaped density we see on the A-capsid portal vertex (Fig. 5D) may therefore act as a docking platform for copies of pUL25 that would be added in the course of a wild type infection and α-helices contributed by pUL36 at a later stage. This platform, however, is evidently insufficient to retain DNA in the capsid, as the absence of pUL25 alone is enough to completely prevent the emergence of propagation-competent, DNA-filled C-capsids (36), underscoring pUL25’s functional versatility as a DNA retention protein and a key player in nuclear egress as well (49).

### Conclusion

Our routine for processing cryo-electron tomograms enabled us to visualize the portal in procapsids in comparison with mature A-capsids. Particularly revealing is the interactions of scaffold with the portal likely to be involved in procapsid assembly. Quantitation of the number of portal vertices per capsid further solidifies a role in assembly. We also gained insight into the intricate maturation-induced changes in the portal vertex, visualizing an outward movement of the portal exceeding vertex movement and thus stabilizing the interaction with the capsid shell.

## Material and Methods

### Cell handling and propagation of virus stocks

Stocks of the UL25 deletion mutant (UL25-null) (36) were generated by using a complementary African Green Monkey cell line (both gifts from Prof. Fred L. Homa, University of Pittsburgh, Pittsburgh, PA). At roughly 75% confluency, monolayers in 175 cm^2^ flasks were infected with virus stock at a MOI of 0.1 in 1 mL of PBS without calcium or magnesium per flask for 40 min at room temperature, followed by 5 min at 37°C. The cells were then overlaid with 15 mL virus growth medium (1x MEM Alpha, Corning, 1.5% (*v/v*) penicillin/streptomycin) and incubated at 37°C. At 48 h post infection, virus-containing medium was clarified by centrifugation at 1,500 rpm for 5 min, and crude cell debris was removed from the supernatant by pelleting at 6,000 rpm for 15 min in a Beckman SW28 rotor. Virus was pelleted for 45 min at 19,000 rpm in a Beckman SW28 rotor. The virus pellets were resuspended in small volumes of PBS without calcium or magnesium, and aliquots of 500 μL were frozen at −80°C. The virus titer was assessed in complementary cells 48 hr. after infection.

Stocks of the procapsid-producing M100 line (35) were generated similarly using the complementary F-3 cell line. Stock titers were assessed as described earlier (5).

### Production of capsids

Roller bottles of African Green Monkey cells were grown to about 75% confluency and infected with UL25-null virus stocks at a MOI of 10. 14 hr post infection, cells had started to round up, but not detach from the surface. Cells were harvested by centrifugation at 1,500 rpm for 5 min, and any virus still attached to the cell surface was removed by incubation with TNE (20 mM Tris, pH 7.4, 500 mM NaCl, 1 mM EDTA) on ice for 15 min. Cells were again pelleted for 5 min at 2,000 rpm, resuspended in TNE, and broken up by sonication. 1.5% (*v/v*) TX-100 was added to solubilize nuclear membranes, followed by pelleting of crude cell debris for 30 min at 6,000 rpm in a Beckman SW28 rotor. The resulting supernatant was supplemented with a cushion of 30% (*w/v*) sucrose in TNE, and a mix of A- and B-capsids was pelleted for 1 h at 20,000 rpm in a Beckman SW28 rotor. The pellets were resuspended in TNE, supplemented with complete protease inhibitor cocktail (Roche), and applied to a preparative gradient containing a range of 20-50% (*w/v*) sucrose in TNE. The gradient was run for 1 h at 22,000 rpm in a Beckman SW55Ti rotor. Distinct bands containing the different capsid species were isolated, pelleted for 50 min at 22,000 rpm in a Beckman SW55Ti rotor, and resuspended in small amounts of TNE to reach an appropriate concentration for cryoET.

Procapsids of the M100 mutant in the viral protease were produced as described earlier (5). They were precipitated with the 6F10 antibody (raised against the triplex protein pUL19) for purification and capsid stability purposes (4).

### Grid preparation and cryo-electron tomography

Purified capsids were mixed with Aurion Anionic 10 nm BSA gold tracers (EMS) at a 1:1 ratio, and 3 μL were applied to Quantifoil copper grids covered with 300-mesh holey carbon (EMS), which were previously plasma-cleaned for 12 s with an argon-oxygen mixture (25% oxygen) in a Model 1020 plasma cleaner (Fischione, Export, PA). Using a Leica EM GP plunger (Leica Microsystems, Buffalo Grove, IL), excess liquid was blotted off for 2.5 s, and grids were flash-frozen in liquid ethane. For the A-capsids, 21 tilt series ranging from −57° to +57° at 3° increments were collected at 20,000x magnification and a defocus target of −2.5 μm on a JEOL JEM-2200FS cryo-electron microscope (JEOL USA, Peabody, MA) operating at 200 kV and equipped with an in-column energy filter and a K2 Summit detector (Gatan, Warrendale, PA). For procapsids, 80 tilt series ranging from −57° to +57° at 3° increments at 81,000x magnification and a −2 μm defocus target were acquired on a 300 kV Titan Krios (Thermo Fisher Scientific, FEI, Hillsboro, OR) equipped with a post-column Model 967 GIF Quantum-LS with a K2 Summit detector (Gatan, Warrendale, PA). In both cases, data were acquired using SerialEM (50).

### Tomogram reconstruction and identification of the portal-bearing vertex

Image processing and tomogram reconstruction were carried out using Bsoft (50, 51). Tomograms were binned 2-fold, resulting in a final pixel size of 3.66 Å for A-capsids and 3.61 Å for procapsids. For both capsids, promising particles were extracted in 450 x 450 x 450 pixel boxes (program bpick) and aligned (program bfind) to a reference of a HSV-1 C-capsid (38) and using an appropriate frequency space mask to compensate for the missing wedge (created with program bmissing). Given the correct alignment parameters, the program bpick was used to extract the 12 vertices for each capsid (150 x 150 x 150 pixel boxes) offset 140 pixels outward from the center of the particle on the fivefold symmetry axes. Using the new Bsoft program, bico, for each set of 12 vertices was cross-correlation against an average calculated from all vertices, and the vertex with the lowest correlation was deemed the portal-bearing vertex. As for the whole capsid alignment, a frequency space mask was used in the correlation calculation (created with program bmissing). In addition, a real space mask was used to isolate a cylindrical region occupied by the penton at the vertex. The mask was created with program beditimg specifying a cylinder centered at 75,75,45 with radius 20 and height 90, and a soft edge with a gaussian width of 3 pixels. The softness of this mask was tuned to avoid high frequency correlations in the comparison of vertex volumes. To address the symmetry mismatch between the fivefold vertex and the twelvefold portal, the best 5-fold orientations of the portal-bearing vertices were found and refined (program bico). The coordinates corresponding to the portal vertices were then transferred to the parameter file referencing the full capsid volumes (program bico). Finally, the average of the whole capsids was calculated with the portal vertex located at the top of the volume. Graphics were prepared using PyMOL (52) and UCSF Chimera (53).

## Data Availability

Maps for the UL25-Null A-capsid and the procapsid, both showing the portal vertex, were deposited in the Electron Microscopy Database under accession numbers EMDB-22378 and EMDB-22379, respectively.

## Acknowledgements

We thank Dr. Fred L. Homa (University of Pittsburgh) for providing us with the HSV-1 UL25-Null mutants and the complementary, UL25-expressing African Green Monkey cell line as well as a wild type African Green Monkey cell line. We thank Dr. Jay Brown for useful discussions. This work was supported by the Intramural Research Program of the National Institute for Arthritis, Musculoskeletal and Skin Diseases, NIH and used the NIH Multi-Institute CryoEM Facility (MICEF).

## Supplementary material

**Fig. S1.**
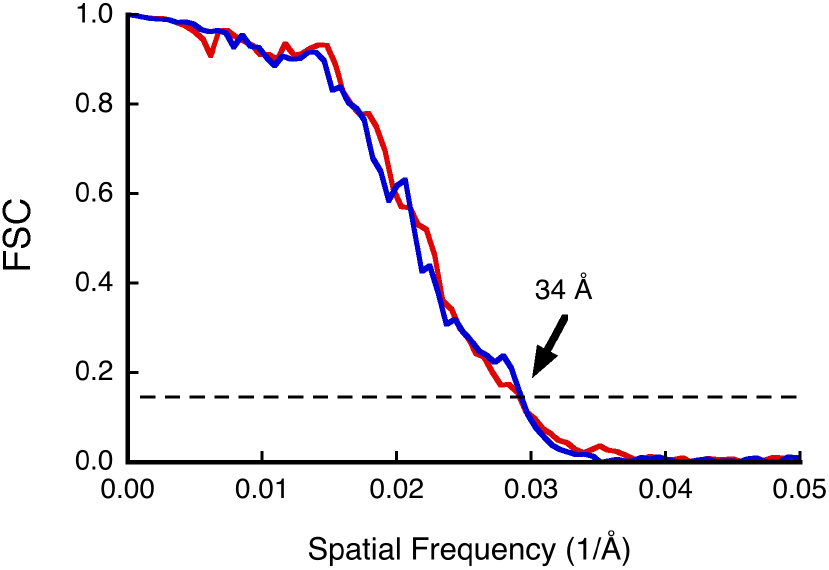
Fourier shell correlation of subtomogram half-set averages of procapsids (red) and A-capsids (blue) with the resolution indicated at a threshold of 0.143 (dashed line). A soft mask covering the capsid shell and portal vertex was used in the analysis.

